# CRISPR-GRIT: Guide-RNAs with Integrated Repair Templates Enable Precise Multiplexed Genome Editing in the Diploid Fungal Pathogen *Candida albicans*

**DOI:** 10.1101/2023.11.27.568926

**Authors:** Christopher J. Cotter, Cong T. Trinh

## Abstract

*Candida albicans* is an opportunistic fungal pathogen that causes severe infections in immunocompromised individuals. Overuse of antifungals coupled with climate change has led to the rapid emergence of antifungal resistance. Thus, there is an urgent need to understand fungal pathogen genetics to develop new antifungal strategies. Genetic manipulation of *C. albicans* is encumbered by its diploid chromosomes which require editing both alleles for elucidating gene function. Even though recent development of CRISPR-Cas systems has facilitated genome editing in *C. albicans*, large-scale functional genomic studies are still hindered by the necessity of co-transforming repair templates for homozygous knockouts. Here, we present CRISPR-GRIT, a repair template-integrated guide RNA design for expedited gene knockouts in *C. albicans,* and demonstrate its utility for multiplexed editing. We envision that this method can be employed for high-throughput library screens and identification of synthetic lethal pairs in both *C. albicans* and other diploid organisms with strong homologous recombination machinery.

## Introduction

*Candida* species are opportunistic yeast that in healthy individuals exists as common commensal members of the microbiome. Microbiome perturbations due to antibiotic use, immune system impairments, and alterations to skin and mucous membrane integrity can lead to *Candida* overgrowth and/or tissue invasion causing potentially life-threatening infections^1^. The prevalence of candidiasis has increased over the past few decades as has the prevalence of antifungal resistance in part due to climate change and overuse of current antifungals^2, 3^. Azole resistance is common among strains of *C. krusei* and *C. glabrata* while echinocandin and polyene resistance is less frequent in *Candida* spp. but has been identified in *C. albicans* and *C. glabrata*^3^*. C. auris* has recently gained significant attention due to its high incidence of antifungal resistance with 91% of isolates being fluconazole-resistant, 12% being amphotericin B resistant, and 12% being echinocandin resistant globally^4^. Additionally, many *C. auris* isolates are multidrug resistant with some being resistant to up to four major classes of antifungal drugs^5^. The alarming increase in antifungal resistance and prevalence of fungal infections requires new strategies for combating fungal pathogens. To rationally design next-generation antifungals, we must first better understand fungal pathogen genetics to appropriately choose drug targets and minimize the probability of evolved resistance.

Of the *Candida* spp., *C. albicans* is the most common cause of candidiasis and is also the most well-studied. Its genetic similarities to the model yeast *Saccharomyces cerevisiae* and relatedness to other *Candida* spp. have made it a popular model fungal pathogen. However, as a diploid, parasexual yeast with no plasmid system, genetic manipulation of *C. albicans* has traditionally been tedious^6^. Conventional approaches to studying functional genomics in *C. albicans* and other diploid or polyploid organisms involve iteratively integrating and recycling selection markers at the desired loci to achieve homozygous knockouts (Fig. 1A). This process is both time-consuming and low throughput.

**Figure 1.**
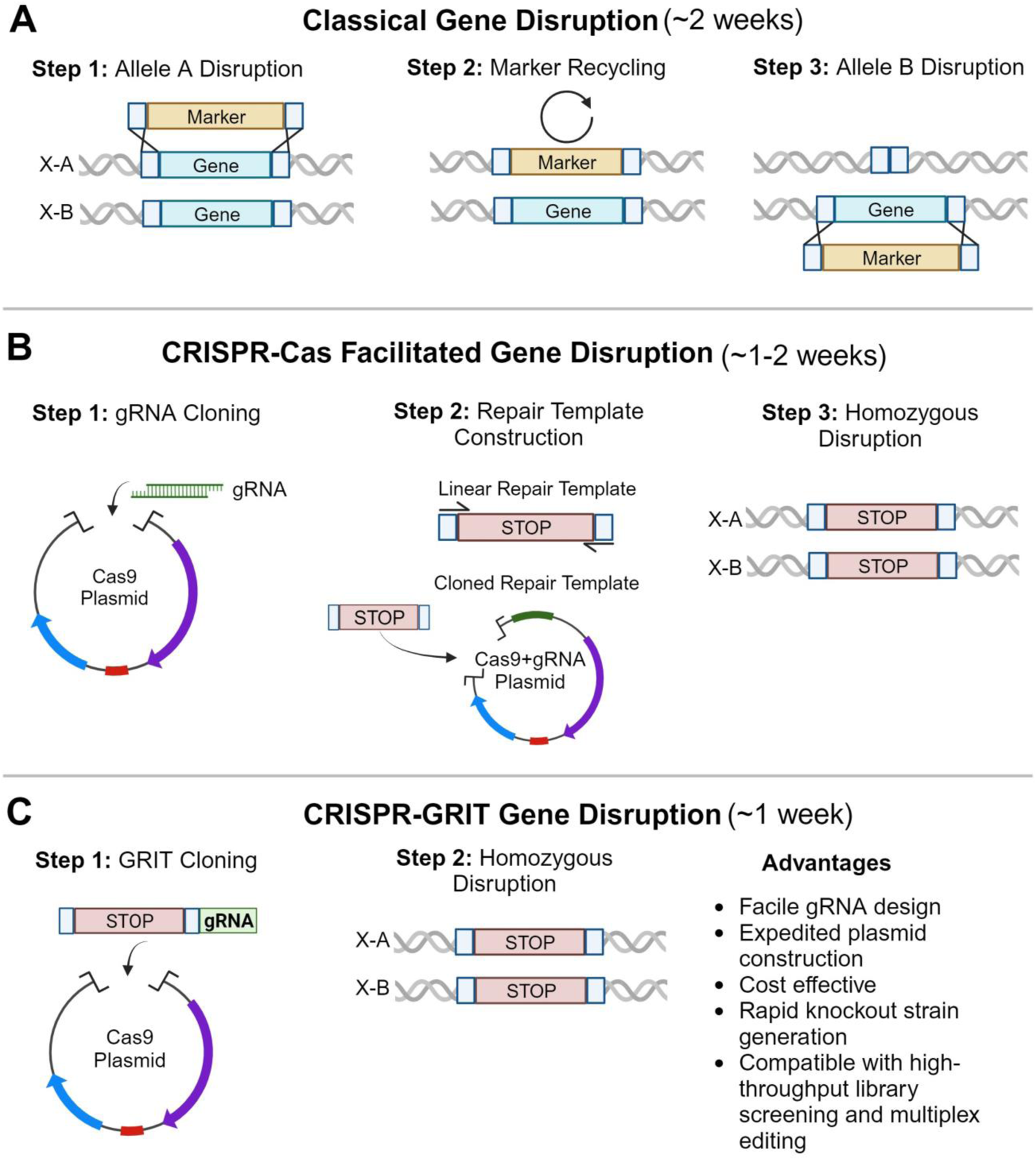
Overview of gene disruption strategies used in *C. albicans*. **(A)** Classical genetics approach involving integration and recycling of selection markers at the locus of interest. **(B)** CRISPR-Cas facilitated gene disruptions. **(C)** CRISPR-GRIT gene disruptions.

The development of CRISPR-Cas systems for *C. albicans* has greatly facilitated genetic studies by streamlining the generation of homozygous knockout strains (Fig. 1B) ^6, 7^. CRISPR-Cas systems utilize a guide RNA (gRNA) and a Cas endonuclease to target and cleave a gene of interest. The induction of DNA double-strand breaks (DSBs) at the target locus activates the host DNA repair machinery allowing for the target gene to be disrupted either inaccurately by introducing indels through non-homologous end joining (NHEJ) or precisely utilizing homologous DNA sequences to introduce desired mutations through homology-directed repair (HDR). In diploid and polyploid organisms, CRISPR-Cas systems can induce DSBs at each allele providing selective pressure for homozygous gene disruption, often eliminating the need for iterative editing. *C. albicans* is HDR dominant and rarely resorts to using NHEJ to repair DNA damage^8^. Therefore, CRISPR-Cas editing in *C. albicans* requires exogenous repair templates with desired mutations to be co-transformed for gene disruption. However, this requirement of repair template co-transformation renders higher throughput CRISPR-Cas gRNA library screens and multiplexed gene editing implausible since each unique gRNA and corresponding repair template must localize to the same cell. With larger numbers of gRNAs and gene targets, the probability of repair template and gRNA co-localization diminishes. Thus, an innovative design method is urgently needed to overcome this current limitation of CRISPR-Cas systems to enable library screens and multiplexed gene editing for diploid or multiploidy organisms such as *C. albicans*.

Currently, large-scale functional genomic studies rely on existing disruption libraries. The GRACE library, based on the disruption of one allele and conditional expression of the second, is the gold standard for elucidating *C. albicans* gene function at scale with conditional knockout strains covering 3,193 of the 6,198 total ORFs^9^. While this resource has been invaluable, there remain close to 3,000 conditional knockout strains to be constructed for complete genome coverage which is a considerable effort. Additionally, this library does not allow for genetic interaction studies or identification of synthetic lethal pairs as only single gene disruptions are available. Efforts to study genetic interactions in *C. albicans* have recently been made possible by CRISPR-Cas gene drives and the discovery of mating competent, stable haploid cells^10^. In these studies, single-gene knockout strains are generated using CRISPR-Cas gene drives in haploid *C. albicans* cells and then mated to generate diploid double-knockout strains. While this CRISPR gene drive technology in haploid cells has enabled the generation of combinatorial deletion mutants, the process for large-scale haploid strain construction and mating still requires significant effort.

In this study we present CRISPR-GRIT, a gRNA with integrated repair template design that fuses the repair template to the gRNA for consolidated cloning, inherent gRNA and repair template co-localization, and efficient CRISPR-Cas editing in *C. albicans* (Fig. 1C). We establish that the CRISPR-Cas editing efficiency can be further improved by the addition of a flanking tRNA^Ala^ at the 5’ end of the CRISPR-GRIT gRNA repair template. Finally, we demonstrate that the CRISPR-GRIT design with discrete transcriptional units can be utilized for efficient multiplexed gene knockouts in *C. albicans.* We anticipate the CRISPR-GRIT design and multiplex knockout capability can enable high-throughput library screens and genetic interaction studies for the identification of synthetic lethal pairs and novel drug targets.

## Results and Discussion

### Design and Implementation of CRISPR-GRIT in C. albicans

Inspired by a homology-integrated guide RNA design utilized in *S. cerevisiae*^11^, we hypothesized that a similar system would be functional in *C. albicans* and eliminate the need for co-transforming linear repair templates. To test this hypothesis, we designed a CRISPR-GRIT gRNA consisting of 50 bp homology arms up and downstream of the gRNA cut site with two premature stop codons and a frameshift mutation that would destroy the PAM fused to the 5’ end of a 20 bp *ADE2* targeting gRNA (Fig. 2A-B). Homozygous disruption of *ADE2* results in a distinct red colony phenotype for rapid knockout screening. This fragment was inserted into the BsmBI stuffer site of the Cas9-gRNA solo plasmid, pV1093, and the resulting plasmid was linearized and transformed into *C. albicans* for integration at the *ENO1* locus. Consistent with previous reports, no red *ADE2* knockouts were observed in the absence of a repair template (Fig. 2D). With the CRISPR-GRIT system, we obtained ∼60% red colonies indicative of homozygous *ADE2* knockouts (Fig. 2E). Additionally, we noticed the total number of colonies per plate was higher with the CRISPR-GRIT system than the conventional CRISPR-Cas design without a repair template, mainly because CRISPR-Cas-induced DNA damage can be lethal in fungi^12^. Even though the CRISPR-GRIT repair template knocks out the conditionally essential gene *ADE2*, integration of the repair template alleviates the stress of DNA damage, suggesting that DNA damage may be more lethal than functional knockouts.

**Figure 2.**
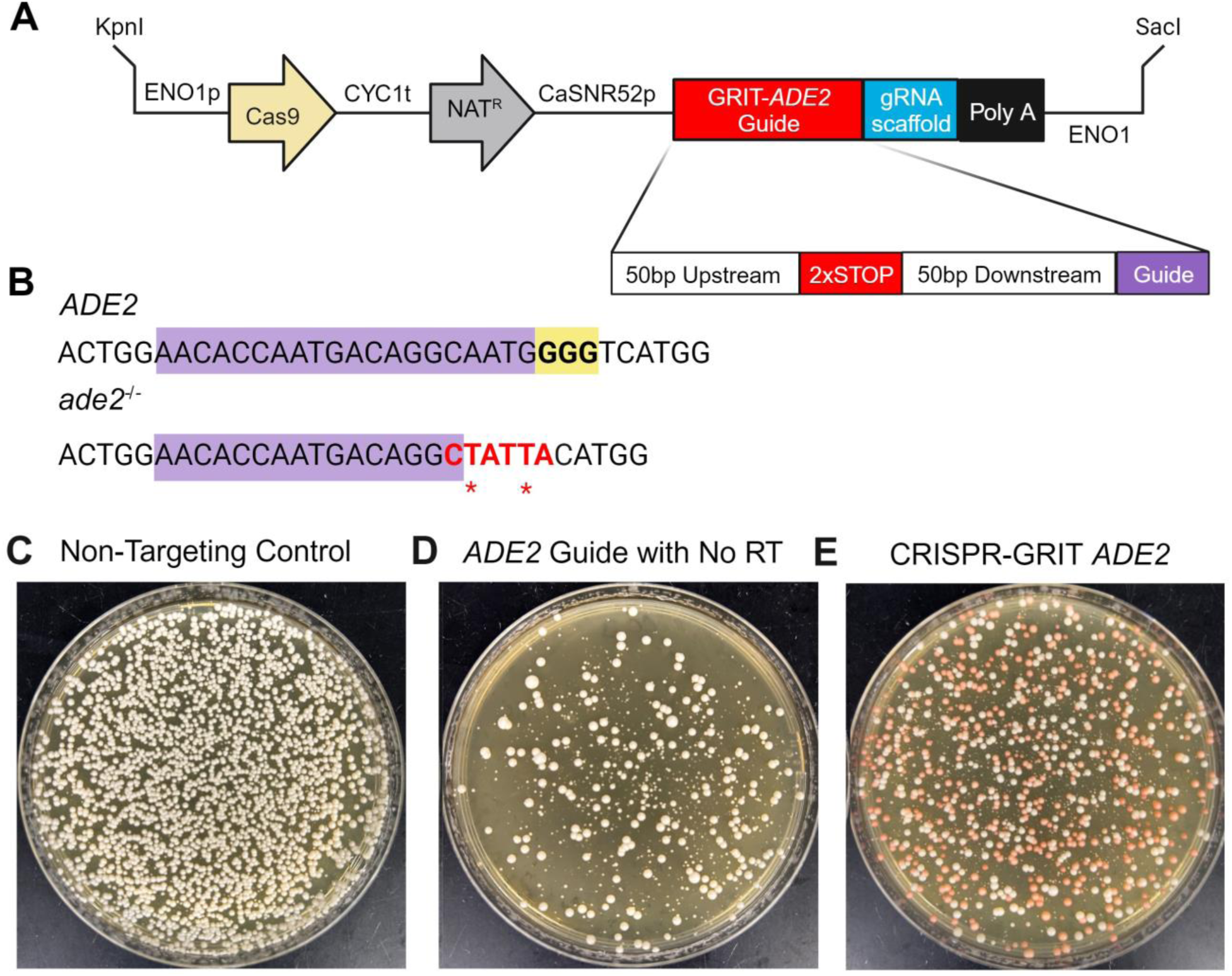
Design and validation of CRISPR-GRIT in *C. albicans.* **(A)** CRISPR-GRIT *ADE2* targeting gRNA and insertion cassette in Cas9-gRNA solo plasmid, pV1093. **(B)** Sequence of the wildtype *ADE2* and CRISPR-GRIT edited *ade2^-/-^* alleles around gRNA cut site. The 20 bp gRNA region and PAM are highlighted in purple and yellow respectively. The premature stop codons introduced by the CRISPR-GRIT repair template are colored in red. **(C-E)** Plate pictures of *C. albicans* transformed with 1 µg of linearized DNA. **(C)** Non-targeting Cas9 control, pV1093, **(D)** pV1093 with an *ADE2* targeting gRNA and no repair template, and **(E)** pV1093 containing the CRISPR-GRIT *ADE2* guide.

### Optimization of CRISPR-GRIT Editing in C. albicans

With a working system established, we then turned to optimizing the CRISPR-GRIT system. We first asked if the homology arms could be shortened as this would significantly reduce the cost of CRISPR-GRIT gRNA synthesis. We varied the homology arm lengths from 50 bp down to 0 in 10 bp increments and evaluated the editing efficiency of each (Fig. 3A). The maximum editing efficiency was observed with 50 bp homology arms (Fig. 3B). We were able to obtain *ADE2* knockouts with homology arm lengths down to 30 bp, albeit with a decrease in editing efficiency with shorter homology arms (Fig 3B). Below 30 bp, no edited colonies were observed (Fig 3B). We chose to continue our optimization using 50 bp homology arms to retain maximal editing efficiency.

**Figure 3.**
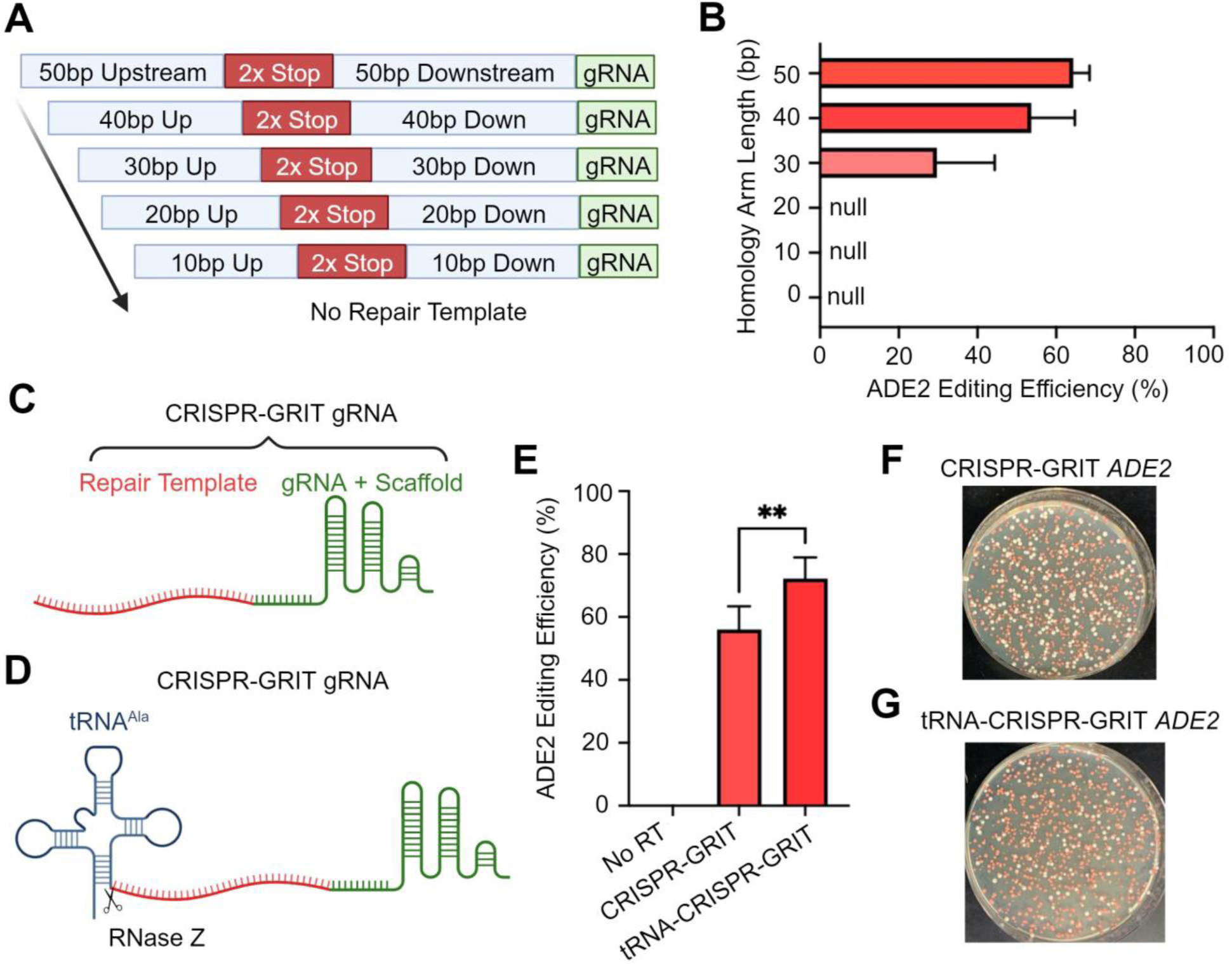
Optimization of CRISPR-GRIT Editing in *C. albicans*. **(A)** Schematic of decreasing CRISPR-GRIT repair template homology arms. **(B)** *ADE2* editing efficiency of CRISPR-GRIT with decreasing repair template homology arm lengths. **(C)** Standard CRISPR-GRIT design. **(D)** CRISPR-GRIT with a flanking 5’ tRNA^Ala^. **(E)** *ADE2* editing efficiency with no repair template, the standard CRISPR-GRIT design, and with a flanking 5’ tRNA^Ala^. **(F)** Plate picture of *C. albicans* transformed with the standard CRISPR-GRIT *ADE2* targeting cassette. **(G)** Plate picture of *C. albicans* transformed with the tRNA-CRISPR-GRIT *ADE2* targeting cassette.

To improve the CRISPR-GRIT editing efficiency and facilitate multiplexed editing, we then investigated the effect of adding a tRNA^Ala^ to the 5’ end of the CRISPR-GRIT gRNA (Fig. 3C-D). Reports have demonstrated that the addition of a flanking tRNA^Ala^ at the 5’ end of the gRNA may improve Cas9 editing in *C. albicans*^13^. Additionally, gRNA-tRNA arrays have been utilized for efficient gRNA multiplexing in *S. cerevisiae* and was a multiplexing design that we intended to test ^14^. Upon the addition of a flanking tRNA^Ala^ to the 5’ end of the CRISPR-GRIT *ADE2* targeting guide, we observed a 29% increase in editing efficiency compared to that lacking the tRNA, improving the editing efficiency from 56% to 72% (Fig 3E-G). This significant increase may be due to increased gRNA stability or nuclear retention^13^.

### CRISPR-GRIT Enables Multiplex Gene Editing in C. albicans

Finally, we investigated the ability to perform multiplexed gene knockouts using the CRISPR-GRIT system. We first attempted to use a gRNA-tRNA array approach where gRNAs separated by tRNAs are expressed on a single transcript and then processed into functional guides by endogenous tRNA-processing enzymes RNase P and RNase Z (Fig. 4A)^14^. We chose to first target *ADE2* and *URA3* since screening for each homozygous knockout phenotype is relatively simple. Homozygous *ADE2* knockouts have a red colony phenotype, homozygous *URA3* knockouts can grow on complete media, but not media lacking uracil, and multiplexed *ADE2/URA3* knockouts should grow and turn red on complete media and not grow on media lacking uracil.

**Figure 4.**
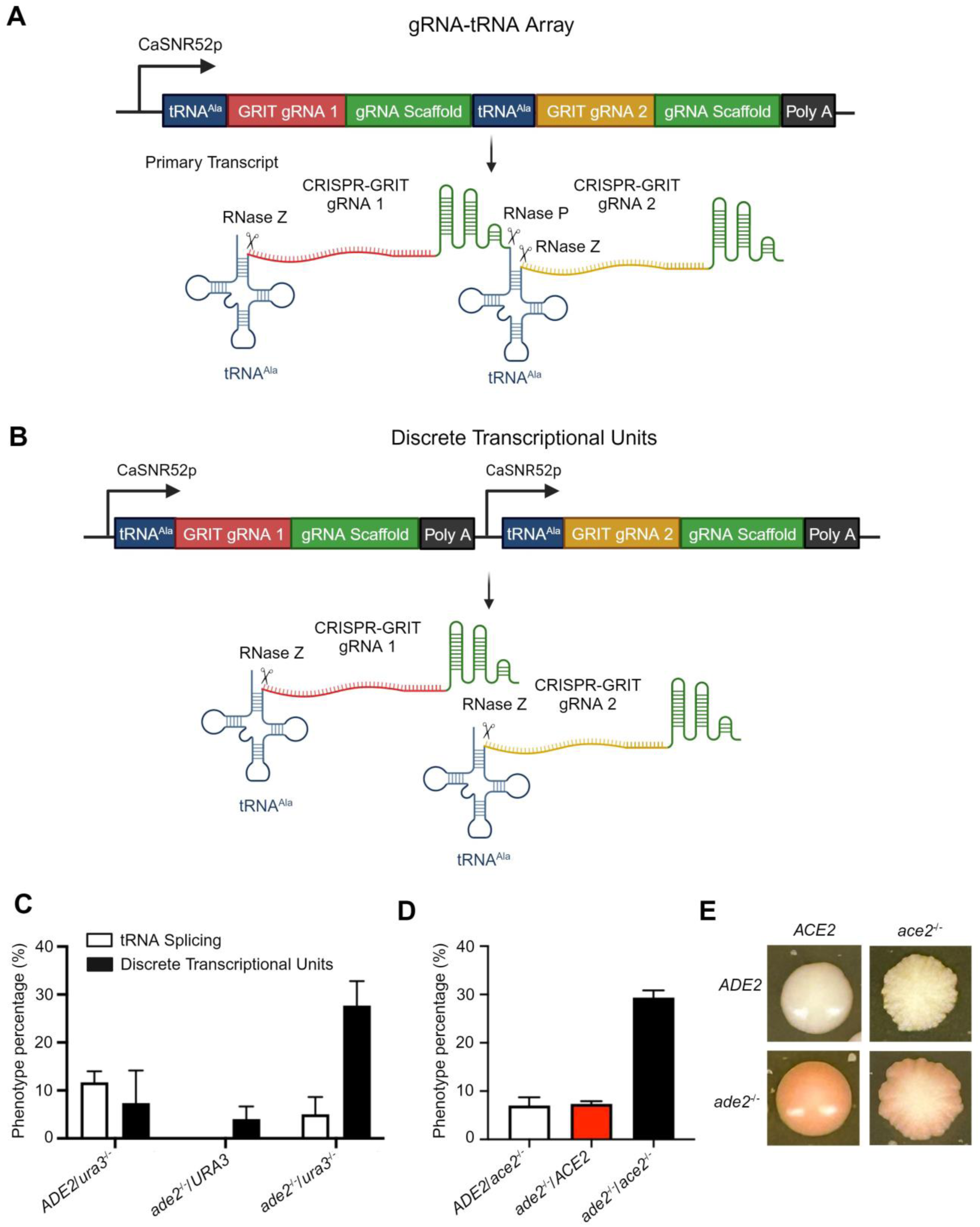
CRISPR-GRIT Multiplexed Gene Editing in *C. albicans*. **(A-B)** DNA and primary transcript of CRISPR-GRIT gRNA-tRNA array **(A)** and discrete transcriptional unit **(B)** multiplexing design. **(C)** *ADE2-URA3* multiplexed editing efficiency of the gRNA-tRNA array and discrete transcriptional unit multiplexing designs. **(D)** *ADE2-ACE2* multiplexed editing efficiency using discrete transcriptional units. **(E)** *C. albicans* colony images of wildtype, *ade2^-/-^, ace2^-/-^,* and *ade2^-/-^/ace2^-/-^* colonies.

Before multiplexing, we first verified that the single *URA3* CRISPR-GRIT gRNA was functional by transforming *C. albicans* with the linearized CRISPR-GRIT *URA3* plasmid and spotting the resulting transformants on rich media and synthetic media lacking uracil (SI Fig. S1). We found that large CRISPR-GRIT *URA3* colonies were typically non-edited escapee cells, whereas smaller colonies had the desired *ura3^-/-^* phenotype and exhibited a growth lag due to the disruption of uracil biosynthesis (SI Fig. S1).

Following validation of the *URA3* CRISPR-GRIT gRNA, we next constructed the *ADE2-URA3* gRNA-tRNA array with the CRISPR-GRIT gRNAs and transformed the resulting linearized plasmid into *C. albicans.* Transformants were spotted on synthetic media with and without uracil and after a 6-day incubation a multiplexed editing efficiency of only 5% was observed (Fig. 4C). Surprisingly, 11.7% of the colonies spotted had an *ADE2/ura3^-/-^* phenotype whereas none had an *ade2^-/-^/URA3* phenotype (Fig. 4C).

We hypothesized that the long repair templates of the CRISPR-GRIT design fused to the gRNA may be interfering with the transcript secondary structure preventing endogenous RNA processing and possibly disrupting gRNA complexation with Cas9. We modeled the gRNA-tRNA array transcripts using RNAfold^15^ and observed a breakdown in tRNA and gRNA scaffold secondary structure upon the addition of the repair templates for the CRISPR-GRIT multiplexing design (SI Fig. 2). Rnase P and Z, responsible for 5’ and 3’ end cleavage respectively, recognize conserved tRNA secondary structures rather than specific RNA sequences, therefore secondary structure alterations may impair recognition and cleavage activity ^16^. Additionally, Rnase Z has poor activity towards tRNAs with long 5’ extensions which the CRISPR-GRIT design adds to the central tRNA in the gRNA-tRNA array ^17^. Inadequate transcript processing could lead to unequal amounts of mature gRNAs or gRNAs with diminished activity leading to the poor multiplexed editing efficiency seen using the gRNA-tRNA array approach.

To avoid transcript secondary structure issues, we opted for a gRNA multiplex design using discrete transcriptional units where each CRISPR-GRIT gRNA was placed under a separate SNR52 promoter (Fig. 4B). After transforming the *ADE2-URA3* targeting CRISPR-GRIT cassette, transformants were again spotted on synthetic media with and without uracil and incubated for 6 days. Using the discrete transcriptional unit approach, we observed an improved multiplexed editing efficiency of 27.7%, whereas 7% of the colonies had an *ADE2/ura3^-/-^* phenotype and 4% had an *ade2^-/-^/URA3* phenotype (Fig. 4C).

While we were able to obtain multiplexed edits, it took a total of 9 days (3 days to obtain transformants, and 6-day incubation after spotting on selective media) to observe the multiplexed *ade2^-/-^/ura3^-/-^*phenotype. Since both genes targeted are conditionally essential and disruption causes a growth lag, we asked whether targeting *ADE2* with a non-essential gene would allow us to observe edits faster or if the kinetics of editing multiple loci was significantly slower than editing a single locus. We chose to target *ADE2* and *ACE2* since *ACE2* is a non-essential gene and mutants have a distinct wrinkly colony phenotype (Fig 4E). We expected that the multiplexed colony phenotype should be red and wrinkled and that multiplexed edited cells should appear after 3 days like that of the single mutants. Indeed after 3 days, multiplexed transformants were observed with a multiplexed editing efficiency of 29.3% demonstrating that multiplexed edits can be obtained rapidly without the need for prolonged incubations (Fig. 4D).

## Conclusions

In this study, we developed a repair template integrated gRNA system, CRISPR-GRIT, for simplified and rapid gene knockouts in the diploid fungal pathogen *C. albicans*. We optimized this system by including a tRNA^Ala^ at the 5’ end of the CRISPR-GRIT gRNA to improve the editing efficiency and demonstrate that edits can be obtained with homology arms as small as 30 bp. We then utilized a discrete transcriptional unit multiplexing design to show that this system can be deployed for efficient multiplex gene knockouts. Large-scale CRISPR-Cas library screens in *C. albicans* have previously been challenging due to the inability to ensure repair template and gRNA co-localization to the same cell. The CRISPR-GRIT design overcomes this barrier by tethering each unique repair template to its associated gRNA, theoretically enabling high-throughput gRNA library screens. Additionally, intrinsic limitations of genetic manipulation in *C. albicans* due to the lack of a plasmid system, diploid chromosomes, and strong homologous recombination machinery impede efforts to perform multiplexed gene editing. The CRISPR-GRIT design is advantageous for multiplexed editing since it applies selective pressure for repair template integration at each allele via chromosomal damage enabling homozygous knockouts. Alternative systems utilized in haploid yeast for multiplexed editing such as base editors lack selective pressure for desired edits and in diploid and polyploid organisms, edits may be outcompeted by homologous recombination of additional copies of wildtype alleles. The multiplexed editing capabilities of CRISPR-GRIT in *C. albicans* open avenues for identifying synthetic lethal pairs and studying genetic interactions at large scales. Further improvements to the system such as Cas9 and gRNA promoter optimization, increased Cas copy number through iterative integration, or utilization of Cas enzymes with increased kinetics or turnover may enhance the editing efficiency and the utility of the CRISPR-GRIT design.

## Materials and Methods

### Strains and Culturing Conditions

*E. coli* NEB DH10β was used for cloning and cultured in LB media at 37°C. *C. albicans* SC5314 (ATCC MYA-2876) was routinely cultured on YPD agar plates or YPD broth at 30°C. *C. albicans URA3* mutants were verified by spotting on synthetic media agar plates without uracil (6.7 g/L yeast nitrogen base without amino acids, 1.92 g/L synthetic dropout medium without uracil 20 g/L glucose, 20 g/L agar).

### Plasmid and GRIT-gRNA Construction

All plasmids and primers are listed in Tables S1-2. The *C. albicans* codon-optimized Cas9 and sgRNA plasmid, pV1093, was a gift from Gerald Fink (Addgene plasmid #111428; http://n2t.net/addgene:111428; RRID: Addgene_111428). Guide RNAs targeting *ADE2* (C3_04520C_A), *URA3* (C3_01350C_A), and *ACE2* (CR_07440W_A) were designed using CASPER^18^. CRISPR-GRIT guide cassettes were designed using a custom Python script. The 20 bp *ADE2* gRNA was constructed by annealing CC-75 and CC-76, oligos containing the *ADE2* seed sequence with overhangs compatible with the BsmbI stuffer of pV1093. To construct CRISPR-GRIT guide RNA plasmids, forward and reverse oligos were combined in a 50 µL PCR to create a homology-integrated guide RNA fragment with flanking BsmBI sites. The resulting fragments were purified using the Omega Biotek E.Z.N.A. Cycle Pure Kit (SKU: D6492-01) and inserted into pV1093-based plasmids via Golden Gate Assembly with BsmBI.

For the construction of the pV1093 plasmid containing the 5’ flanking tRNA^Ala^, pV1093-CatRNA, a gene strand CatRNABsmBISacffold was synthesized from Eurofins containing homology to SNR52 promoter and downstream of the pV1093 gRNA scaffold region. The SNR52 promoter was amplified from pV1093 using primers CC-371 and CC-372. pV1093 was digested with NotI and SacII, and the PCR-amplified SNR52 promoter and CatRNABsmBIScaffold gene strand were inserted into the digested backbone via Gibson Assembly.

For gRNA-tRNA multiplexing, the first gRNA was PCR amplified using primers CC-377 and CC-378 while the second was PCR amplified using primers CC-379 and CC-380. The PCR-amplified gRNAs were assembled into pV1093-CatRNA via Golden Gate Assembly with BsmBI.

For discrete transcriptional unit gRNA multiplexing, a destination plasmid lacking the guide RNA scaffold, pV1093-Empty, was first constructed the same as pV1093-CatRNA, but using a separate gene strand, CaBsmBIStuffer, instead of CatRNABsmBIScaffold. The first gRNA was PCR amplified using primers CC-377 and CC-463 and the second gRNA was PCR amplified using primers CC-464 and CC-380. The PCR-amplified gRNAs were assembled into pV1093-Empty via Golden Gate Assembly with BsmBI.

Correct plasmids were confirmed by colony PCR using primers CC-79/CC-22 and Sanger sequencing using primer CC-79 for single gRNAs or gRNA-tRNA multiplexed gRNAs or CC-466 and CC-467 for discrete transcriptional unit multiplexed gRNAs. Before transformation into *C. albicans*, 5-10 µg of each plasmid was linearized using KpnI-HF and SacI-HF in an overnight digest at 37°C. The digests were purified using the Omega Biotek E.Z.N.A. Cycle Pure Kit (SKU: D6492-01) and the total purified products were used for subsequent transformations.

### Transformation of C. albicans

The protocol developed by Walther and Wendland was used as the standard lithium acetate transformation method with the following modifications ^19^. A freshly streaked *C. albicans* colony was used to inoculate in 10 mL of minimal media (6.7 g/L yeast nitrogen base without amino acids, 5 g/L glucose, 0.36 g/L potassium acetate) for 12-16 hours at 30°C, 250 rpm. The culture was then diluted to an OD600 of 0.3 in 25 mL of minimal media and incubated at 30°C, 250 rpm for 5-6 hours until the culture reached an OD600 of ∼1.8. Cells were collected by centrifugation at 900 x g for 5 minutes, washed once with 12 mL of sterile water, and resuspended to an OD600 of 30 in 100 mM lithium acetate titrated to a pH of 5 using glacial acetic acid. 100 µL of cells were aliquoted in sterile 1.5 mL tubes. 1 ug of linearized DNA, 10 µL of 10 mg/mL herring sperm DNA, and 600 µL of 50% PEG 3350 dissolved in 100 mM lithium acetate were then added to each tube. Transformation mixtures were incubated for 18-20 hours static at 30°C then heat shocked at 44°C for 15 minutes. Cells were then pelleted at 900 x g for 5 minutes and washed with 1 mL of YPD. Cells were resuspended in 2 mL of YPD in a 15 mL culture tube and recovered for 4 hours at 30°C shaking at 250 rpm before plating serial dilutions on YPD agar plates supplemented with 250 µg/mL nourseothricin for selection. Plates were incubated at 30°C for 3 days before counting colonies.

## ACKNOWLEDGEMENTS

This research is financially supported in part by the ORII Seed Fund at The University of Tennessee, Knoxville, the DARPA YFA award and Director Fellowship (D17AP00023), and the DOE BER Genomic Science Program (DE-SC0019412). The views, opinions, and/or findings contained in this article are those of the authors and should not be interpreted as representing the official views or policies, either expressed or implied, of the funding agencies. The mention of trade names or commercial products in this publication is solely for the purpose of providing specific information and does not imply a recommendation or endorsement by the funding agencies.

## AUTHOR CONTRIBUTIONS

Conceptualization: CT; Investigation: CC; Methodology: CC; Formal Analysis: CC, CT; Visualization: CC, CT; Funding Acquisition: CT; Project Administration: CT; Supervision: CT; Writing-Review & Editing: CC, CT.

## CONFLICT OF INTEREST

The authors declare no competing financial interest.

## Notes

### Competing Interest Statement

The authors have declared no competing interest.

## References

1. Lopes, J. P.; Lionakis, M. S. Pathogenesis and virulence of *Candida albicans*. Virulence 2022, 13 (1), 89–121. DOI: 10.1080/21505594.2021.2019950.

2. Bongomin, F.; Gago, S.; Oladele, R. O.; Denning, D. W. Global and Multi-National Prevalence of Fungal Diseases-Estimate Precision. J Fungi (Basel) 2017, 3 (4). DOI: 10.3390/jof3040057. Nnadi, N. E.; Carter, D. A. Climate change and the emergence of fungal pathogens. PLoS Pathog 2021, 17 (4), e1009503. DOI: 10.1371/journal.ppat.1009503.

3. Perlin, D. S.; Rautemaa-Richardson, R.; Alastruey-Izquierdo, A. The global problem of antifungal resistance: prevalence, mechanisms, and management. Lancet Infect Dis 2017, 17 (12), e383–e392. DOI: 10.1016/S1473-3099(17)30316-X.

4. Chen, J.; Tian, S.; Han, X.; Chu, Y.; Wang, Q.; Zhou, B.; Shang, H. Is the superbug fungus really so scary? A systematic review and meta-analysis of global epidemiology and mortality of *Candida auris*. BMC Infectious Diseases 2020, 20 (1), 827. DOI: 10.1186/s12879-020-05543-0.

5. Jacobs, S. E.; Jacobs, J. L.; Dennis, E. K.; Taimur, S.; Rana, M.; Patel, D.; Gitman, M.; Patel, G.; Schaefer, S.; Iyer, K.;, et al. Candida auris Pan-Drug-Resistant to Four Classes of Antifungal Agents. Antimicrobial Agents and Chemotherapy 2022, 66 (7), e00053–00022. DOI: doi:10.1128/aac.00053-22.

6. Vyas, V. K.; Barrasa, M. I.; Fink, G. R. A *Candida albicans* CRISPR system permits genetic engineering of essential genes and gene families. Science Advances 2015, 1 (3), e1500248. DOI: 10.1126/sciadv.1500248.

7. Wensing, L.; Sharma, J.; Uthayakumar, D.; Proteau, Y.; Chavez, A.; Shapiro, R. S. A CRISPR Interference Platform for Efficient Genetic Repression in *Candida albicans*. mSphere 2019, 4 (1), 10.1128/msphere.00002-00019. DOI: doi:10.1128/msphere.00002-19.

8. Vyas, V. K.; Bushkin, G. G.; Bernstein, D. A.; Getz, M. A.; Sewastianik, M.; Barrasa, M. I.; Bartel, D. P.; Fink, G. R. New CRISPR Mutagenesis Strategies Reveal Variation in Repair Mechanisms among Fungi. mSphere 2018, 3 (2). DOI: 10.1128/mSphere.00154-18. Yao, S.; Feng, Y.; Zhang, Y.; Feng, J. DNA damage checkpoint and repair: From the budding yeast *Saccharomyces cerevisiae* to the pathogenic fungus Candida albicans. Comput Struct Biotechnol J 2021, 19, 6343–6354. DOI: 10.1016/j.csbj.2021.11.033.

9. Skrzypek, M. S.; Binkley, J.; Binkley, G.; Miyasato, S. R.; Simison, M.; Sherlock, G. The Candida Genome Database (CGD): incorporation of Assembly 22, systematic identifiers and visualization of high throughput sequencing data. Nucleic Acids Res 2017, 45 (D1), D592–d596. DOI: 10.1093/nar/gkw924. Roemer, T.; Jiang, B.; Davison, J.; Ketela, T.; Veillette, K.; Breton, A.; Tandia, F.; Linteau, A.; Sillaots, S.; Marta, C.; et al. Large-scale essential gene identification in Candida albicans and applications to antifungal drug discovery. Mol Microbiol 2003, 50 (1), 167–181. DOI: 10.1046/j.1365-2958.2003.03697.x. Fu, C.; Zhang, X.; Veri, A. O.; Iyer, K. R.; Lash, E.; Xue, A.; Yan, H.; Revie, N. M.; Wong, C.; Lin, Z.-Y.; et al. Leveraging machine learning essentiality predictions and chemogenomic interactions to identify antifungal targets. Nature Communications 2021, 12 (1), 6497. DOI: 10.1038/s41467-021-26850-3.

10. Shapiro, R. S.; Chavez, A.; Porter, C. B. M.; Hamblin, M.; Kaas, C. S.; DiCarlo, J. E.; Zeng, G.; Xu, X.; Revtovich, A. V.; Kirienko, N. V.;, et al. A CRISPR-Cas9-based gene drive platform for genetic interaction analysis in *Candida albicans*. Nat Microbiol 2018, 3 (1), 73–82. DOI: 10.1038/s41564-017-0043-0.

11. Bao, Z.; Xiao, H.; Liang, J.; Zhang, L.; Xiong, X.; Sun, N.; Si, T.; Zhao, H. Homology-Integrated CRISPR–Cas (HI-CRISPR) System for One-Step Multigene Disruption in *Saccharomyces cerevisiae*. ACS Synthetic Biology 2015, 4 (5), 585–594. DOI: 10.1021/sb500255k.

12. Mendoza, B. J.; Zheng, X.; Clements, J. C.; Cotter, C.; Trinh, C. T. Potency of CRISPR-Cas Antifungals Is Enhanced by Cotargeting DNA Repair and Growth Regulatory Machinery at the Genetic Level. ACS Infectious Diseases 2023. DOI: 10.1021/acsinfecdis.3c00342.

13. Ng, H.; Dean, N. Dramatic Improvement of CRISPR/Cas9 Editing in *Candida albicans* by Increased Single Guide RNA Expression. mSphere 2017, 2 (2), 10.1128/msphere.00385-00316. DOI: doi:10.1128/msphere.00385-16.

14. Zhang, Y.; Wang, J.; Wang, Z.; Zhang, Y.; Shi, S.; Nielsen, J.; Liu, Z. A gRNA-tRNA array for CRISPR-Cas9 based rapid multiplexed genome editing in *Saccharomyces cerevisiae*. Nature Communications 2019, 10 (1), 1053. DOI: 10.1038/s41467-019-09005-3.

15. Lorenz, R.; Bernhart, S. H.; Höner zu Siederdissen, C.; Tafer, H.; Flamm, C.; Stadler, P. F.; Hofacker, I. L, ViennaRNA Package 2.0. Algorithms for Molecular Biology 2011, 6 (1), 26. DOI: 10.1186/1748-7188-6-26. Hofacker, I. L. Vienna RNA secondary structure server. Nucleic Acids Research 2003, 31 (13), 3429–3431. DOI: 10.1093/nar/gkg599.

16. Sekulovski, S.; Trowitzsch, S. Transfer RNA processing – from a structural and disease perspective. Biological Chemistry 2022, 403 (8-9), 749–763. DOI: doi:10.1515/hsz-2021-0406.

17. Pellegrini, O.; Nezzar, J.; Marchfelder, A.; Putzer, H.; Condon, C. Endonucleolytic processing of CCA-less tRNA precursors by RNase Z in *Bacillus subtilis*. The EMBO Journal 2003, 22 (17), 4534–4543. DOI: 10.1093/emboj/cdg435.

18. Mendoza, B.; Fry, T.; Dooley, D.; Herman, J.; Trinh, C. T. CASPER: An Integrated Software Platform for Rapid Development of CRISPR Tools. CRISPR J 2022, 5 (4), 609–617. DOI: 10.1089/crispr.2022.0025. Mendoza, B. J.; Trinh, C. T. Enhanced guide-RNA design and targeting analysis for precise CRISPR genome editing of single and consortia of industrially relevant and non-model organisms. Bioinformatics 2018, 34 (1), 16–23. DOI: 10.1093/bioinformatics/btx564.

19. Walther, A.; Wendland, J. An improved transformation protocol for the human fungal pathogen *Candida albicans*. Current Genetics 2003, 42 (6), 339–343. DOI: 10.1007/s00294-002-0349-0.

