## Supplementary material for "CRISPR-GRIT: Guide-RNAs with Integrated Repair Templates Enable Precise Multiplexed Genome Editing in the Diploid Fungal Pathogen *Candida albicans*": Figures S1-2, Tables S1-S2

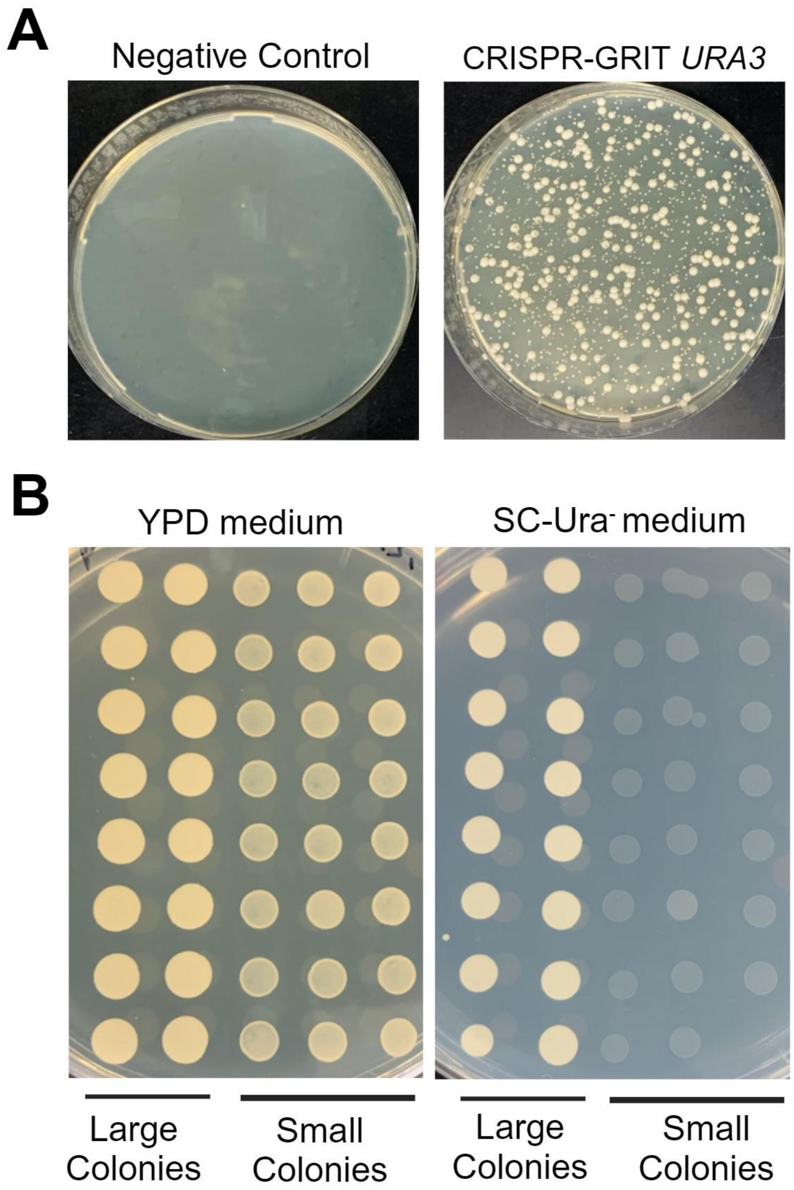


**Figure S1:** Validation of CRISPR-GRIT *URA3* gRNA. **(A)** *C. albicans* transformation with 1 µg CRISPR-GRIT *URA3* plated on YPD+NAT 250 µg/mL. The negative control contained no DNA. **(B)** Large and small colonies from the CRISPR-GRIT *URA3* transformation plate were dissolved in sterile water and spotted on YPD and SC-URA$^-$ plates to screen for *URA3* knockouts. Small colonies from the CRISPR-GRIT *URA3* transformation plate grew on YPD, but not SC-URA$^-$ indicative of a *URA3* knockout. Large colonies grew on both plates suggesting they are cells that have escaped CRISPR-Cas editing.

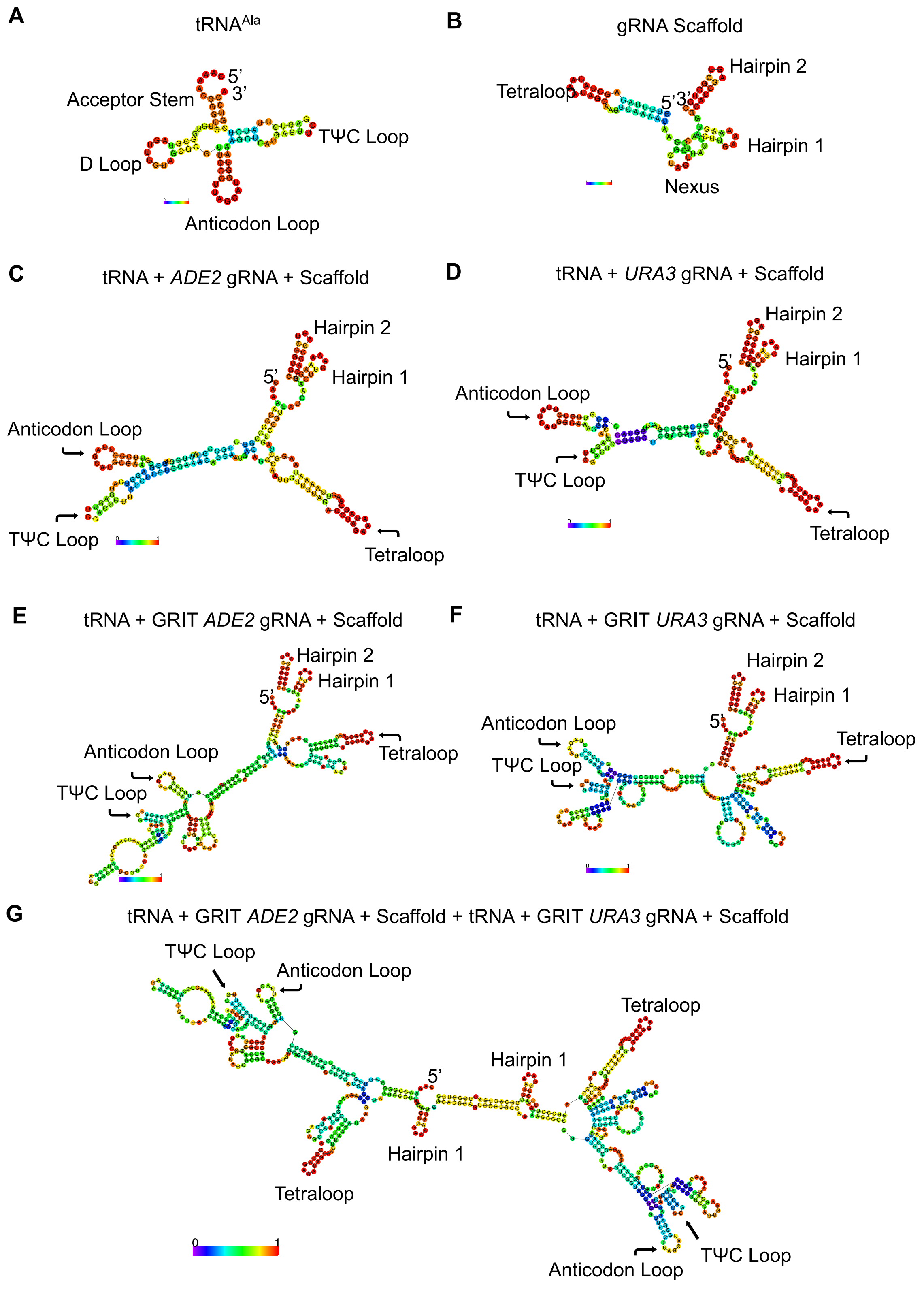
A
tRNA^Ala
5'
3'
Acceptor Stem
TΨC Loop
D Loop
Anticodon Loop
B
gRNA Scaffold
Hairpin 2
Tetraloop
5'3'
Hairpin 1
Nexus
C
tRNA + ADE2 gRNA + Scaffold
Hairpin 2
5'
Hairpin 1
Anticodon Loop
TΨC Loop
Tetraloop
D
tRNA + URA3 gRNA + Scaffold
Hairpin 2
5'
Hairpin 1
Anticodon Loop
TΨC Loop
Tetraloop
E
tRNA + GRIT ADE2 gRNA + Scaffold
Hairpin 2
Hairpin 1
5'
Tetraloop
Anticodon Loop
TΨC Loop
F
tRNA + GRIT URA3 gRNA + Scaffold
Hairpin 2
Hairpin 1
5'
Anticodon Loop
Tetraloop
TΨC Loop
G
tRNA + GRIT ADE2 gRNA + Scaffold + tRNA + GRIT URA3 gRNA + Scaffold
TΨC Loop
Anticodon Loop
Tetraloop
Hairpin 1
5'
Hairpin 1
Tetraloop
0
1
TΨC Loop
Anticodon Loop

**Figure S2:** RNA secondary structure modeling of CRISPR-GRIT transcript elements using RNAfold. **(A)** Secondary structure of the $tRNA^{Ala}$ and notable secondary structure features: Acceptor stem, D loop, Anticodon loop, and TΨC loop. **(B)** Secondary structure of the SpCas9 gRNA scaffold and notable secondary structure features: Tetraloop, Nexus, Hairpin 1, and Hairpin 2. **(C-D)** Secondary structure of $tRNA^{Ala}$ with a single 20 bp gRNA and scaffold targeting **(C)** *ADE2* or **(D)** *URA3*. Upon fusion to the 20 bp gRNA and scaffold, the $tRNA^{Ala}$ D loop is no longer present. **(E-F)** Secondary structure of $tRNA^{Ala}$ with a single GRIT-gRNA and scaffold targeting **(E)** *ADE2* or **(F)** *URA3*. The GRIT-gRNA repair templates add significant secondary structure to the primary transcript. **(G)** Secondary structure of gRNA-tRNA array multiplexed design with GRIT-gRNAs targeting *ADE2* and *URA3*. In addition to the $tRNA^{Ala}$ D loop absence, Hairpin 2 of the gRNA scaffolds is also absent.

**Table S1.** List of plasmids used in this study.

| **Plasmid** | **Strain** | **Source** |
|---|---|---|
| pV1093 | *E. coli* DH10β | Vyas et al. (Addgene #111428)[1] |
| pV1093-CatRNA | *E. coli* DH10β | This Study |
| pV1093-Empty | *E. coli* DH10β | This Study |
| pV1093-*ADE2* | *E. coli* DH10β | This Study |
| CRISPR-GRIT *ADE2* | *E. coli* DH10β | This Study |
| CRISPR-GRIT *ADE2* 40bp HA | *E. coli* DH10β | This Study |
| CRISPR-GRIT *ADE2* 30bp HA | *E. coli* DH10β | This Study |
| CRISPR-GRIT *ADE2* 20bp HA | *E. coli* DH10β | This Study |
| CRISPR-GRIT *ADE2* 10bp HA | *E. coli* DH10β | This Study |
| CRISPR-GRIT *ADE2* 100bp HA | *E. coli* DH10β | This Study |
| CRISPR-GRIT *ADE2* 200bp HA | *E. coli* DH10β | This Study |
| tRNA-CRISPR-GRIT *ADE2* | *E. coli* DH10β | This Study |
| tRNA-CRISPR-GRIT *URA3* | *E. coli* DH10β | This Study |
| tRNA-CRISPR-GRIT *ACE2* | *E. coli* DH10β | This Study |
| GTA-CRISPR-GRIT *ADE2-URA3* | *E. coli* DH10β | This Study |
| DTU-CRISPR-GRIT *ADE2-URA3* | *E. coli* DH10β | This Study |
| DTU-CRISPR-GRIT *ADE2-ACE2* | *E. coli* DH10β | This Study |

**Table S2.** List of primers and gene fragments used in this study.

| Primer/Gene Fragment | Sequence (5' to 3') | Description |
|---|---|---|
| CatRNABsmbIScaffold | CGAGACTTGCGTAAACTATTTTTAATTTGCAAAACGGGCGTGTGGCGTAGTTGGTAGCGCGTTCCCTTAGCATGGGAAAGGTCATGAGTTCGACTCTTATCTCGTCCAGAGACGGAACCTCGCAGACTTTTCCGTCTCGTTTTAGAGCTAGAAATAGCAAGTTAAAATAAGGCTAGTCCGTTATCAACTTGAAAAAGTGGCACCGAGTCGGTGCTTTTTTTCTCGAGTTTTTTTATCGAGTGTTTAAGGATAATGATAACTGAAGAGAAGAATTAGTTTTGCCGCCACCGCGGGTTTGCCTCTGATTAAATAAAAAAAAAGCTGG | tRNA, BsmBI stuffer, and SpCas9 sgRNA scaffold gene fragment for creation of pV1093-CatRNA |
| CaBsmbIStuffer | CGAGACTTGCGTAAACTATTTTTAATTTGGAGACGGAACCTCGCAGACTTTTCCGTCTCTTTTTTTCTCGAGTTTTTTTATCGAGTGTTTAAGGATAATGATAACTGAAGAGAAGAATTAGTTTTGCCGCCACCGCGGGTTTGCCTCTGATTAAATAAAAAAAAAGCTGG | BsmBI stuffer and SpCas9 sgRNA scaffold gene fragment for creation of pV1093-Empty |
| CC-466 | CGTAgtcatgagtcagacttatc | Discrete Transcriptional Unit gRNA #1 sequencing primer |
| CC-467 | agctcagtgattaagagtaaagatg | Discrete Transcriptional Unit gRNA #2 sequencing primer |
| CC-79 | gcaccaattacgtaccaag | Amplification of sgRNA cassette FW and gRNA sequencing primer |
| CC-22 | gttggtggggcaataactcc | Amplification of sgRNA cassette RV |
| CC-377 | AAACGTCTCTATTTGCAAAACGGGCGTGTGG | Multiplexing gRNA #1 FW |
| CC-378 | TTTCGTCTCTTTTGgcaccgactcggtgc | gRNA-tRNA Array Multiplex gRNA #1 RV |

| | | |
|---|---|---|
| CC-379 | AAACGTCTCTCAAAACGGGCGTGTGG | gRNA-tRNA Array Multiplex gRNA #2 FW |
| CC-380 | TTTCGTCTCAAAAAAgcaccgactcggtgc | Multiplexing gRNA #2 RV |
| CC-463 | TTTCGTCTCgcgtagtcatgagtcagacttatcattatccttaaacactcg | Discrete Transcriptional Unit Multiplex gRNA #1 RV primer |
| CC-464 | AAACGTCTCTTACGgtgattagacttagtccgttc | Discrete Transcriptional Unit Multiplex gRNA #2 FW primer |
| CC-371 | attattctctagttttgacggcggccgcagtgattagacttag | SNR52 primer FW |
| CC-372 | caaattaaaaatagtttacgcaagtctcg | SNR52 primer RV |
| CC-75 | atttgAACACCAATGACAGGCAATGg | ADE2 gRNA FW |
| CC-76 | aaaacCATTGCCTGTCATTGGTGTTc | ADE2 gRNA RV |
| HiCRISPRFW | AAACGTCTCTATTTGGAATCAACCCCATCTAATGTAGATCCCTTAACTGGAACACCAATGACAGGCTATTACATGGCCGCCACCATAC | GRIT-ADE2 FW for pV1093 |
| HiCRISPRRV | TTTCGTCTCCAAAACCATTGCCTGTCATTGGTGTTTTGCTGGTGCAGGTGGTGCTGCTCATTTGCCAGGTATGGTGGCGGCCATG | GRIT-ADE2 RV |
| CC-373 | AAACGTCTCTGTCCAGAATCAACCCCATCTAATGTAGATCCCTTAACTGGAACACCAATGACAGGCTATTACATGGCCGCCACCATACCT | GRIT-ACE2 FW for pV1093-CatRNA |
| CC-375 | AAACGTCTCTGTCCATTGAAGGATTAAAACAGGGAGCTAAAGAAACCACCACCAACCAAGAGCCAtaataaATTGATGTTAGCTGAATTA | GRIT-URA3 FW |
| CC-376 | TTTCGTCTCCAAAACTCTTGGCTCTTGGTTGGTGGATTCTCCATATGCTAATGATCCCACTGATGATAATTCAGCTAACATCAATttatt | GRIT-URA3 RV |
| CC-351 | AAACGTCTCTATTTGATCTAATGTAGATCCCTTAACTGGAACACCAATGACAGGCTATTACATGGCCGCCACCATAC | GRIT-ADE2 40bp HA FW |

| | | |
|---|---|---|
| CC-352 | TTTCGTCTCCAAAACCATTGCCTGTCATTGGTGTTAGGTGGTGCTGCTCATTTGCCAGGTATGGTGGCGGCCATG | GRIT-ADE2 40bp HA RV |
| CC-353 | AAACGTCTCTATTTGGATCCCTTAACTGGAACACCAATGACAGGCTATTACATGGCCGCCACCATAC | GRIT-ADE2 30bp HA FW |
| CC-354 | TTTCGTCTCCAAAACCATTGCCTGTCATTGGTGTTGCTCATTTGCCAGGTATGGTGGCGGCCATG | GRIT-ADE2 30bp HA RV |
| CC-355 | AAACGTCTCTATTTGCTGGAACACCAATGACAGGCTATTACATGGCCGCCACCATAC | GRIT-ADE2 20bp HA FW |
| CC-356 | TTTCGTCTCCAAAACCATTGCCTGTCATTGGTGTTCAGGTATGGTGGCGGCCATG | GRIT-ADE2 20bp HA RV |
| CC-357 | AAACGTCTCTATTTGAATGACAGGCTATTACATGGCCGCCAACACCA | GRIT-ADE2 10bp HA FW |
| CC-358 | TTTCGTCTCCAAAACCATTGCCTGTCATTGGTGTTGGCGGCCATG | GRIT-ADE2 10bp HA RV |
| Cal_GRITADE2_100 | CATGGTCCTGCTGGAGTTCGTGAAACGTCTCTATTTGAACAGTTGCAACAGGAATACCTCTTGGCATCTGTACTATAGAGTGTAACGAATCAACCCCATCTAATGTAGATCCCTTAACTGGAACACCAATGACAGGCTATTACATGGCCGCCACCATACCTGGCAAATGAGCAGCACCACCTGCACCAGCAATGATACATTTCAAGCCACGCTTTGGTGCTTCAATAGCATACTCAGACATTAACACCAATGACAGGCAATGGTTTTGGAGACGAAACGTCGCCGTCCAGCTCGACCAGC | GRIT-ADE2 100bp HA |
| Cal_GRITADE2_200 | AAACGTCTCTATTTGTGTTCAACATATATTGGTTCATCTCAGTCAACCATTTTGAGTCATAGGCGCCTAATATTCTGATTGCCAACAATGCAGCATTAGTACTATTGTTAATAGCAACAGTTGCAACAGGAATACCTCTTGGCATCTGTACTATAGAGTGTAACGAATCAACCCCATCTAATGTAGATCCCTTAACTGGAACACCAATGACAGGCTATTACATGGCCGCCACCATACCTGGCAAATGAGCAGCACCACCTGCACCAGCAATGATACATTTCAAGCCACGCTTTGGTGCTTCAATAGCATACTCAGACATTCTATGAGG | GRIT-ADE2 200bp HA |

| | | |
|---|---|---|
| | AGTTCTGTGTGCACTTACGATTGTGAGTTCAAATGGCACACCAAATTGTTTCAAAATACGAGCACCAACTGCCATAACTGGTAGATCCGAATAACACCAATGACAGGCAATGGTTTTGGAGACGAAA | |
| ACE2-FW | AAACGTCTCTGTCCATGCATTTATCACCTTTGAAAAAACAATTACCAAACACTCCCACAAAGCAAtaataaCACCATTGAATGGAGTCC | GRIT-ACE2 FW |
| ACE2-RV | TTTCGTCTCCAAAACCTGTCACCATTGAATGGAGTGTAATGGTTGCTTTGAGTTTGGTGATATAACTGGACTCCATTCAATGGTGttat | GRIT-ACE2 RV |

### References

(1) Vyas, V. K.; Barrasa, M. I.; Fink, G. R. A *Candida albicans* CRISPR system permits genetic engineering of essential genes and gene families. *Science Advances* **2015**, *1* (3), e1500248. DOI: 10.1126/sciadv.1500248.
